# Autoantibodies and anti-microbial antibodies: Homology of the protein sequences of human autoantigens and the microbes with implication of microbial etiology in autoimmune diseases

**DOI:** 10.1101/403519

**Authors:** Peilin Zhang

## Abstract

Autoimmune disease is a group of diverse clinical syndromes with defining autoantibodies within the circulation. The pathogenesis of autoantibodies in autoimmune disease is poorly understood. In this study, human autoantigens in all known autoimmune diseases were examined for the amino acid sequences in comparison to the microbial proteins including bacterial and fungal proteins by searching Genbank protein databases. Homologies between the human autoantigens and the microbial proteins were ranked high, medium, and low based on the default search parameters at the NCBI protein databases. Totally 64 human protein autoantigens important for a variety of autoimmune diseases were examined, and 26 autoantigens were ranked high, 19 ranked medium to bacterial proteins (69%) and 27 ranked high and 16 ranked medium to fungal proteins (66%) in their respective amino acid sequence homologies. There are specific autoantigens highly homologous to specific bacterial or fungal proteins, implying potential pathogenic roles of these microbes in specific autoimmune diseases. The computational examination of the primary amino acid sequences of human autoantigens in comparison to the microbial proteins suggests that the environmental exposure to the commensal or pathogenic microbes is potentially important in pathogenesis of a majority of autoimmune diseases, providing a new direction for further experimental investigation in searching for new diagnostic and therapeutic targets for autoimmune diseases.

## Introduction

Autoimmune disease is characterized by the presence of circulating autoantibodies. Some autoantibodies, for example, anti-smith antibodies, are more specific and pathognomonic to a specific autoimmune disease such as systemic lupus erythematosus (SLE)(1, 2). Others such as anti-nuclear antibody, are non-specific and present in many clinical diseases or even normal healthy individuals (1, 2). The questions of how and why these autoantibodies are generated in patients remain largely unanswered. Microbiome study demonstrated the presence of trillions of various microbes within the body, and there is an intimate symbiotic relationship between these microbes and the human host in various aspects of human tissue and organ functions (3-5). In addition, there are numerous species of microbial DNA within the blood circulation without the culturable microbes (6). The presence of the microbial DNA in blood without culturable microbes raises two possibilities: the blood culture methods are insensitive to detect the microbes but the microbes are present within the circulation (7), or alternatively the intact microbes are destroyed by the host immune system but the microbial DNA and proteins are present in the circulation, eliciting the human immune response to these microbial DNA/proteins. The human immune responses to the microbial DNA and/or proteins may manifest as anti-microbial antibodies with potential cross-reactivity to human tissues through molecular mimicry (8, 9). In the process of identifying the infectious agents in Crohn’s disease and Sjogren’s syndrome, we have identified a group of microbial proteins from the commensal or pathogenic microbes reacting to patients’ plasma, and we showed that there were elevated antibodies within the circulation of patients reacting to the microbial proteins (8). Furthermore, we showed that there is a cross reactivity of anti-microbial antibodies to the human tissues. It is reasonable to assume that the anti-microbial antibodies in response to the microbes, either pathogenic or commensal, can be detected as autoantibodies in autoimmune diseases, given the presence of cross-reactivity of anti-microbial antibodies against human tissues (8).

In current study, I took one step further to examine the primary amino acid sequences of all known human autoantigens in autoimmune diseases and compared these amino acid sequences to the microbial proteins including the bacterial and fungal proteins, using information from the Genbank and BLAST search tools publically available at National Center for Biotechnology Information (NCBI) (https://blast.ncbi.nlm.nih.gov/Blast.cgi). Our results are surprising and more than two third of the human autoantigens important for a variety of autoimmune diseases are homologous to the microbial proteins including bacterial and fungal proteins. These microbial proteins may elicit the human antibody responses with potential cross-reactivity to the human tissues, leading to specific human tissue damage and autoimmune diseases. The presence of anti-microbial antibodies in circulation in autoimmune diseases suggests a potential new mechanism of autoantibody production and autoimmune diseases. The computational analysis of the primary protein sequences for homology represents the initial step toward understanding the production of autoantibodies in autoimmune diseases.

## Methods

All known human autoantigens associated with human autoimmune diseases are listed in Table 1. The protein (amino acid) sequences of these human autoantigens were searched from the Genbank (https://www.ncbi.nlm.nih.gov/protein/?term=), and the specific amino acid sequences and/or specific Genbank accession numbers were used for BLASTP search against the bacterial database (Bacteria (taxid:2)) and fungal database (Fungi (taxid:4751)) using the previously defined search parameters (default parameters). The homology was ranked as low, medium and high using previously defined (default) parameters, and denoted in color from NCBI (blue — low, medium— pink, red — high) (Figure 1). Screenshot photos were taken to demonstrate the homology between the human proteins with the various microbial proteins with colored graphical illustrations and the specific amino acid alignments between the human autoantigens and the perspective microbial proteins (bacterial and/or fungal). The specific homology scores from the default NCBI search algorism including compositional matrix adjustment, positive identity and gaps were not used because of the lack of expertise from the author’s perspective. The low, medium and high homology definitions based on the colored coded graphics were used to estimate the similarity of two protein sequences from two separate species. Unique specific microbes were also denoted in Table 1 for specific homologous human autoantigens. Each and all human autoantigen with their respective search data are illustrated in a book to be published later (in preparation). The current writing is the summary of all the search data with a goal to point to an entirely different direction for further experimental investigation for the mechanism of autoantibodies and autoimmune diseases.

**Figure 1.**
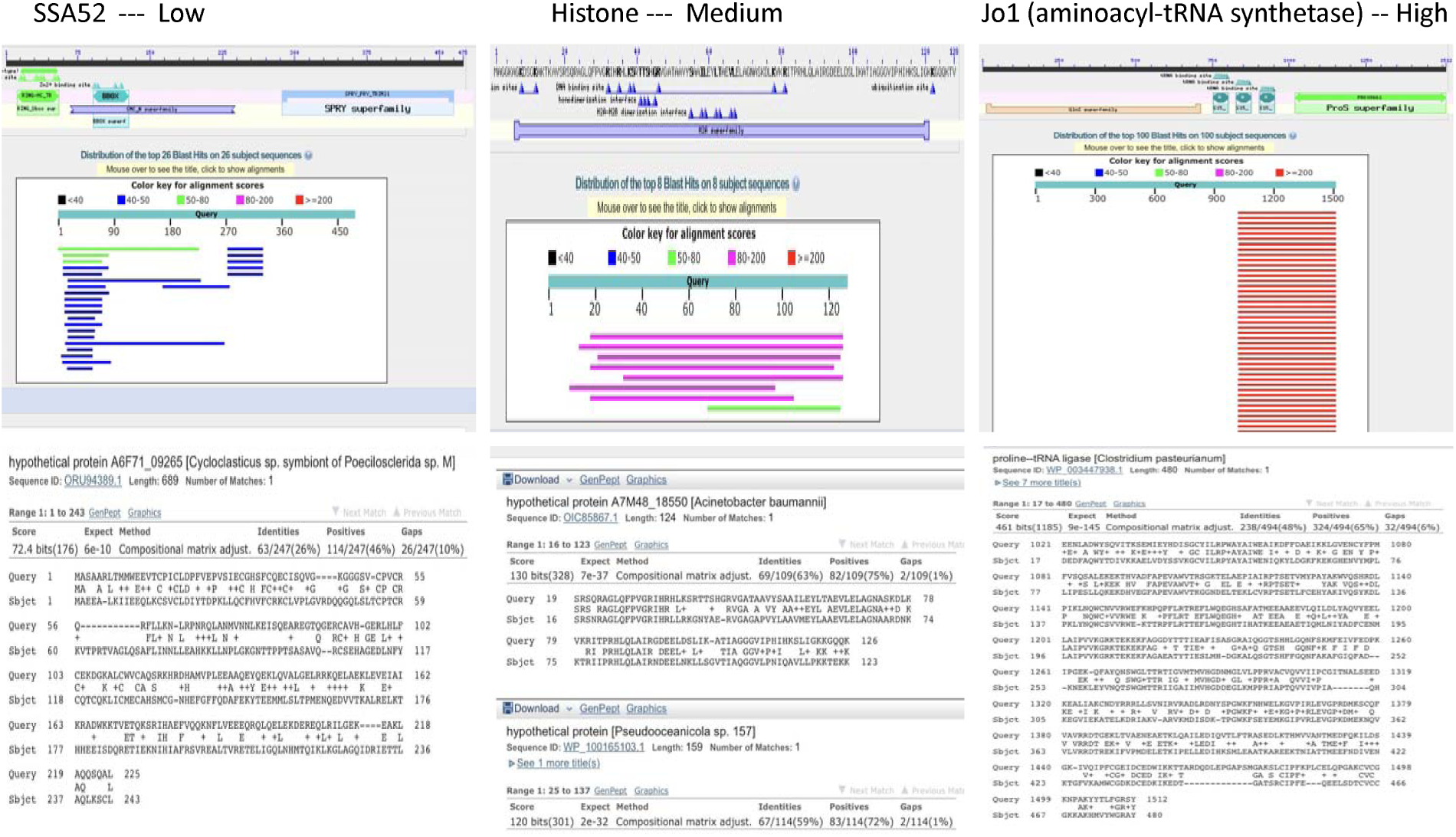
Definition of protein sequence homology with colored graphic illustration and the sequence alignment between the human autoantigens and the microbial proteins. The BLASTP search parameters were of the default setting, and the colored graphical illustrations were from the NCBI.

**Table 1:**
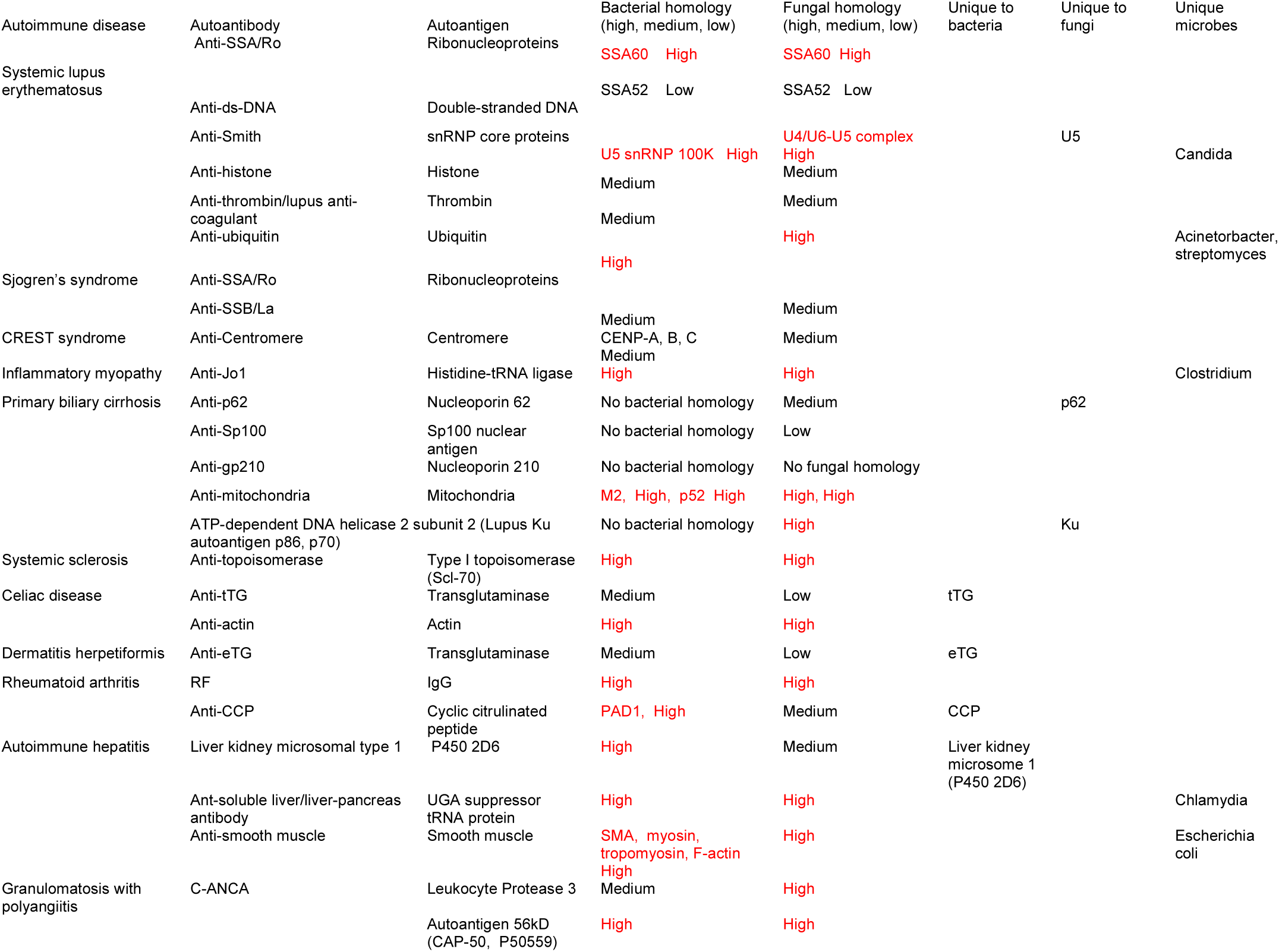
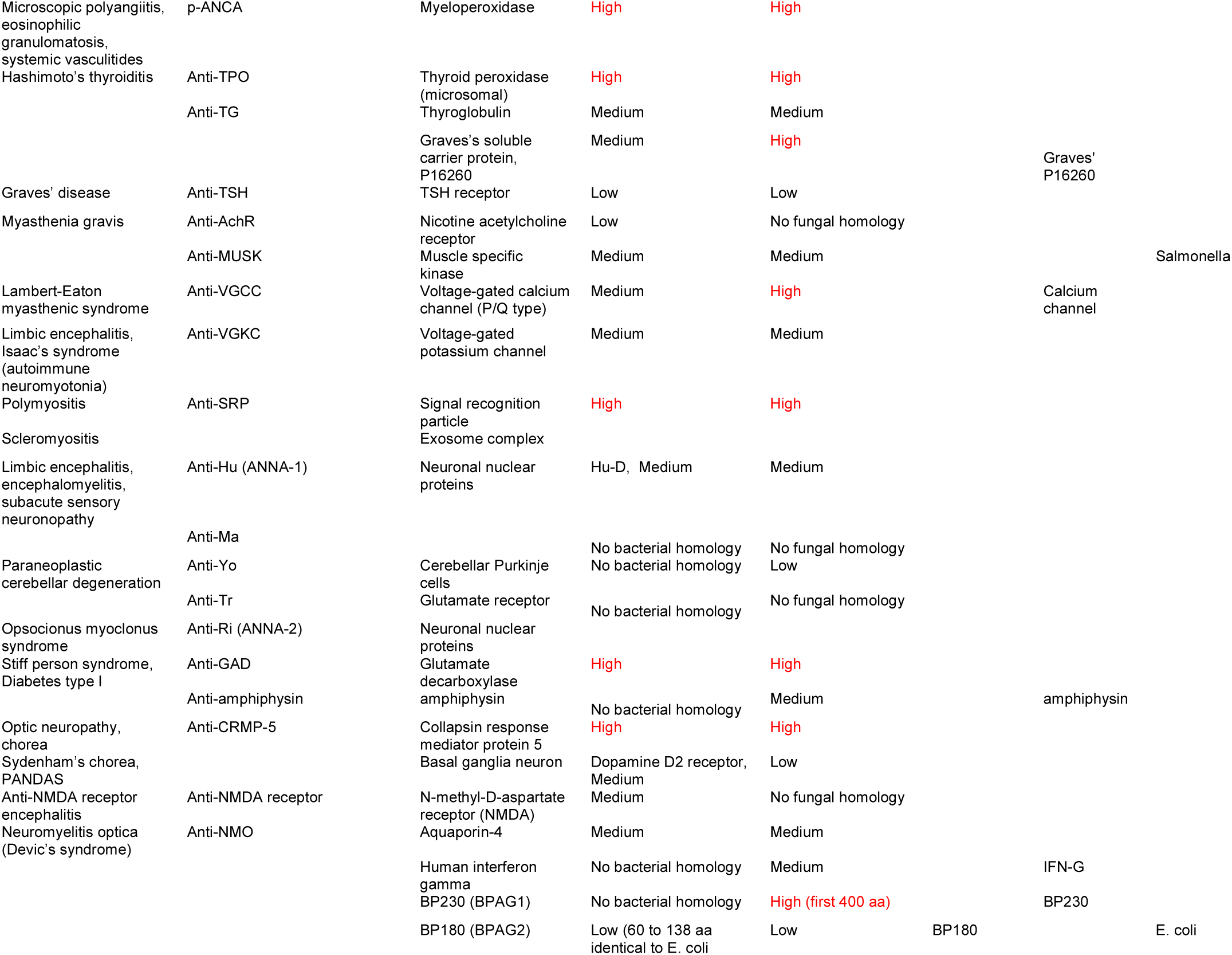
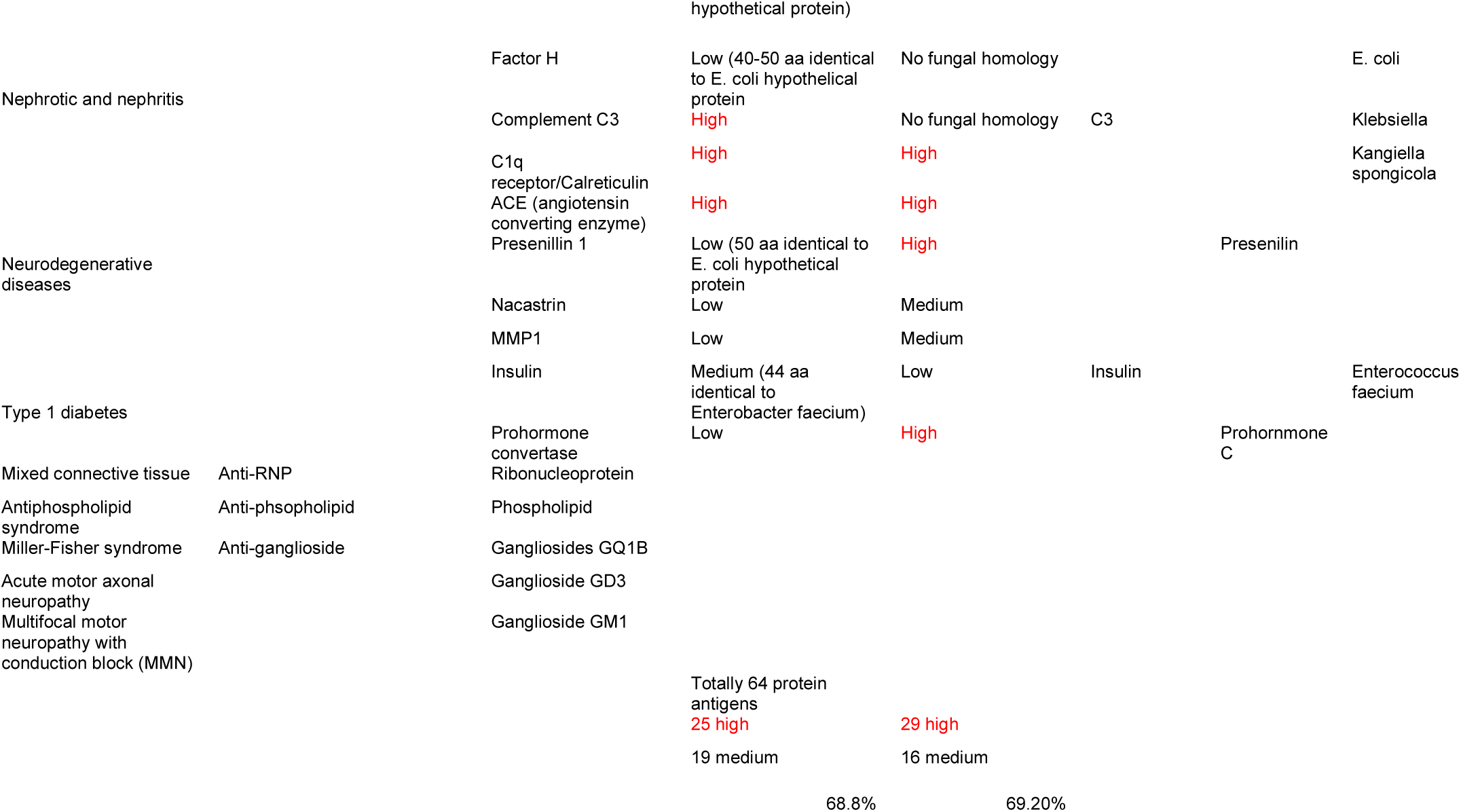
Human autoantigens and homologies to microbial proteins in autoimmune diseases.

## Results

### 1. Genbank search, definition of low, medium and high homology between the human autoantigens and the microbial proteins

There are three examples of BLASTP searches for definitions of low, medium and high homology between the amino acid sequences of human autoantigens and the microbial proteins (Figure 1). Using the Genbank accession number for each individual human autoantigen (SSA52/Ro52, Genbank accession number AAA36581, and Histone H2A, Genbank accession number P20671, anti-Jo1 autoantigen, Genbank accession number P07814) and BLASTP search, three matched results were shown in Figure 1. The low, medium, and high homologies between the SSA52/R052, Histone H2A and Jo1 and the microbial proteins were illustrated by blue, pink and red colored graphics followed by the amino acid sequence alignments with the respective microbial proteins from specific bacterium or fungi. It is worth noting that antibody production requires a small stretch of amino acid (epitope) within an appropriate antigenic structure. The examination of the amino acid sequences can only show the homology and alignment between the human proteins and the microbial proteins without predicting the three-dimensional structure. As a general rule, the experimental significance of this homology search is limited to the knowledge of computational prediction, and the predictive information requires vigorous experimental validation to be clinically relevant. The importance and clinical significance of these human autoantigens in the specific autoimmune diseases is beyond the scope of this study.

### 2. Microbial proteins reactive to human plasmas in Crohn’s disease and Sjogren’s syndrome are highly homologous to human protein homologues

We have previously identified a panel of bacterial proteins reactive to human plasmas from patients with Crohn’s disease and Sjogren’s syndrome, and we demonstrated that there were elevated anti-microbial antibodies within the circulation of these patients (8). We also demonstrated that the specific antibodies against the microbial proteins produced in vitro can cross react to human tissues. Currently, the amino acid sequences of these microbial proteins were used to search Genbank human protein database to compare the microbial proteins with human proteins (Figure 2). This is a reverse exercise of the methods in Figure 1. The query sequences from the bacterial proteins were used to search the human protein database (Homo sapiens (taxid:9606)). The homologies between the microbial proteins EF-G, ATP5a from *Staphylococcus aureus /pseudintermedius (10)*, Hsp65 from *mycobacterium avium subspecies hominissuis/Mycobacterium tuberculosis*, and EF-Tu from *Escherichia coli* to the human proteins were significantly high, and the antibody against the microbes can cross react to the human tissues as shown previously (8). The clinical significance of these microbial proteins and their respective antibodies within the circulation in Crohn’s disease and Sjogren’s syndrome have been previously discussed (8, 9).

**Figure 2.**
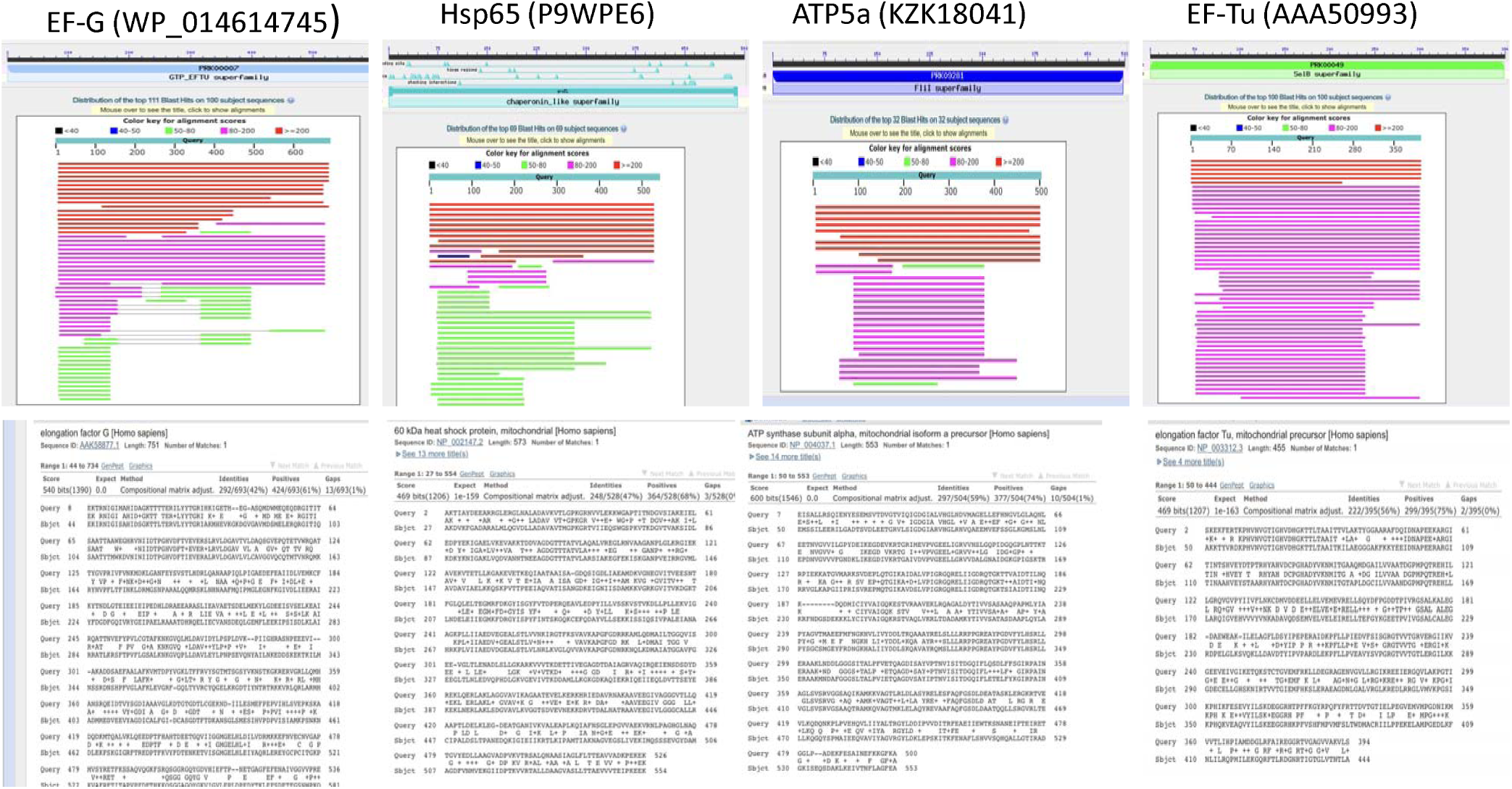
Highly homologous bacterial proteins identified in Crohn’s and Sjogren’s patients to human proteins by the default BLASTP search with graphic illustrations and protein sequence alignments. Individual microbial proteins and their Genbank accession numbers were listed on top of the graphics.

### 3. Homology of all human autoantigens to microbial proteins

Using the same principle and search methods with identical search parameters, the known human autoantigens important for human autoimmune diseases were examined, and the results were listed in Table 1. Totally 64 protein autoantigens in a variety of autoimmune diseases were examined against the microbial protein databases including bacterial database (Bacteria (taxid:2)) and fungal database (Fungi (taxid:4751)). There were 25 autoantigens highly homologous to the bacterial proteins, 29 highly homologous to the fungal proteins. 19 human autoantigens showed medium homology to the bacterial proteins and 16 to fungal proteins. Two examples of high protein sequence homology between the human autoantigens and the microbial proteins (bacterial and fungal) are shown in Figure 3. Combining the high and medium homology groups, there are 68.8% of human autoantigens showing medium to high homology to the bacterial proteins and 69% to the fungal proteins. Phospholipids can also be antigenic in autoimmune diseases such as Guillain-Barre syndrome or anti-phospholipid antibody syndromes, but the phospholipids serve as haptens in antibody response, and the haptens usually need to combine with carrier proteins to be antigenic (11). The phospholipids can derive from plasma membranes of eukaryotic or prokaryotic cells, and it is difficult to determine the sources of these phospholipids in these patient populations. It is also unclear if the patients with anti-phospholipid antibodies carry other circulating anti-microbial antibodies against other microbial proteins.

**Figure 3.**
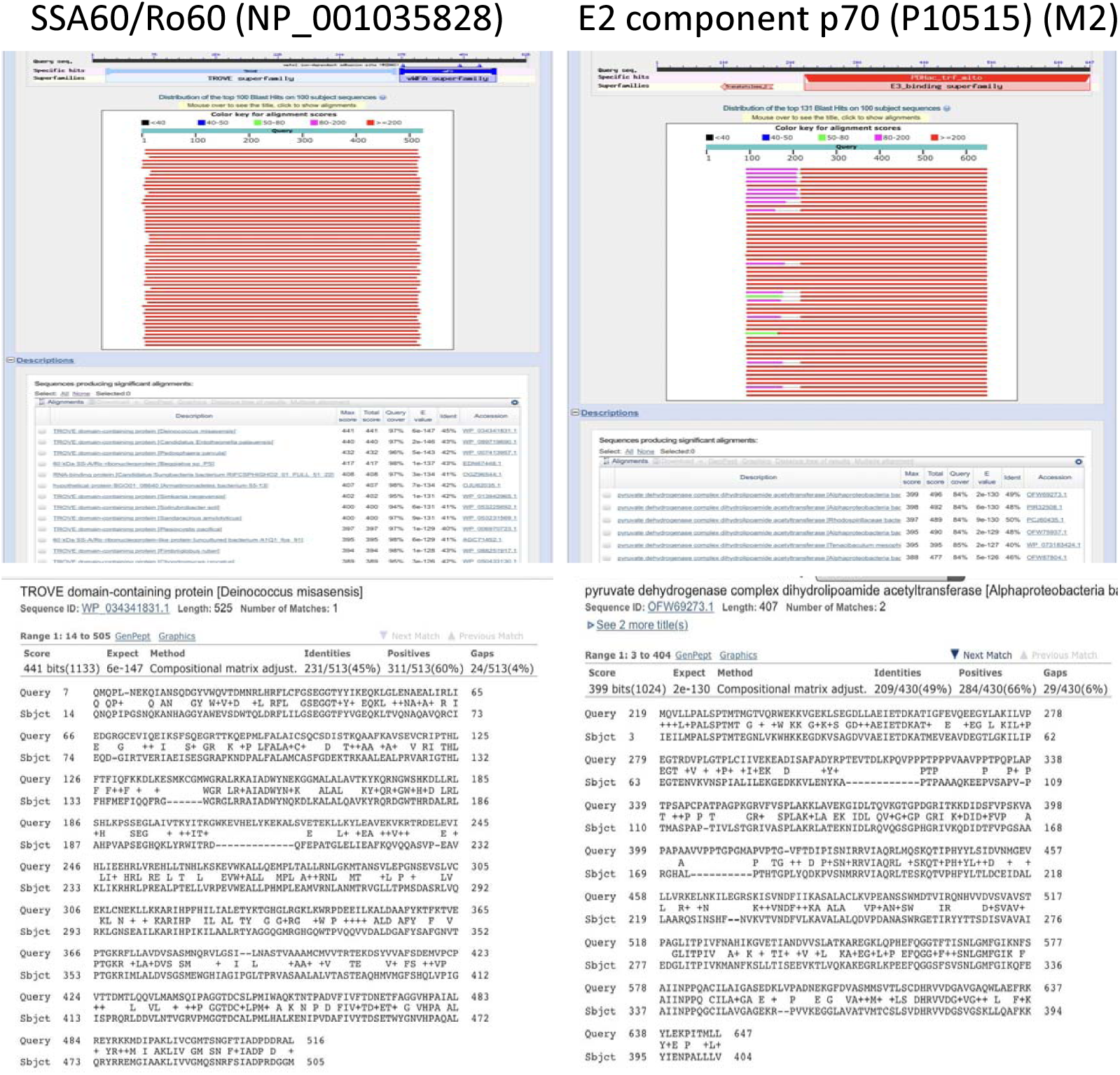
Representative high homology alignments between the human autoantigens and the microbial proteins (SSA60 and mitochondrial protein M2 complex E2 component with their Genbank accession numbers).

There is other important information that is medically relevant to clinical management of a variety of patients with autoimmune diseases, and this information requires experimental validation. Human autoantigen U5 ribonuclear protein in SLE is highly homologous to that of the *Candida albicans*. Human proinsulin sequence is homologous to a specific hypothetical protein of *Enterococcus faecium (12)*. Histidine tRNA ligase of anti-Jo1 antibody important for inflammatory myopathy is highly specific to that of *Clostridium*, and the antigens from anti-smooth muscle antibodies in autoimmune hepatitis were found to be specific to those of *Escherichia coli*. Specific clinical management plan can be devised if these relationships between the specific autoantigens and specific microbial proteins are experimentally validated.

Among the autoantigens highly homologous to the microbial proteins (red colored), most commonly seen were enzymes with catalytic functions in both human host cells and the microbial cells. The structural proteins in cytoskeleton are also common, and these structural proteins are well conserved across the microbial or all species with important functions in cell division and cell mobility (Table 2). Others proteins with various functions such as nucleoproteins, regulatory proteins, immune related proteins and ion channels are also identified. It is conceivable that the human immune responses to these microbial proteins lead to antibody production and these anti-microbial antibodies will cross-react to the human tissue through molecular mimicry, leading to collateral human tissue damage.

**Table 2:**
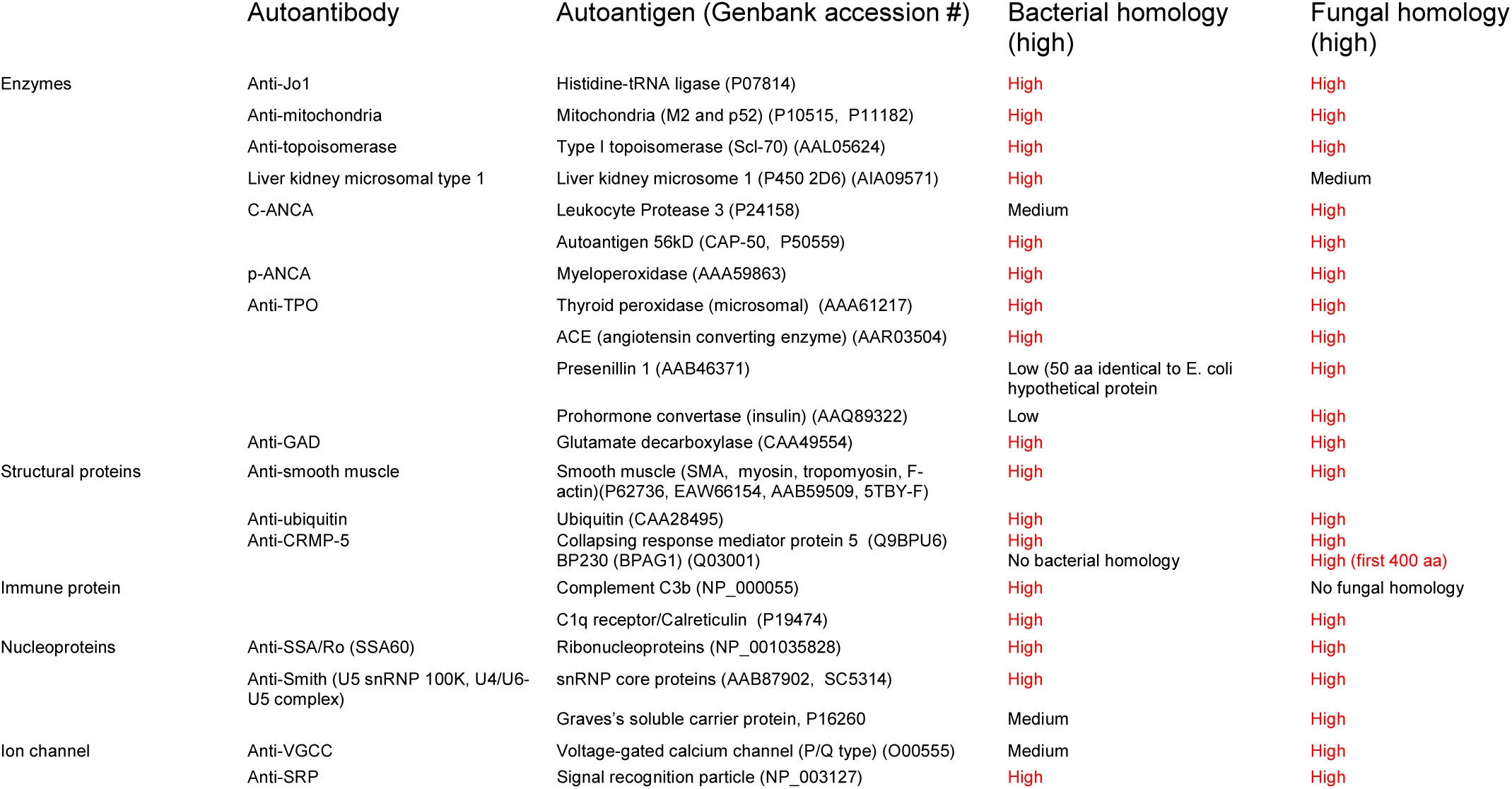
Protein classification of the autoantigens and microbial proteins.

## Discussion

It is well known that the B-cells and plasma cells are critical for autoimmune diseases (13). Through examination of all human autoantigens against the bacterial and fungal protein databases, approximately two third of the human autoantigens are significantly homologous to the microbial proteins including bacterial and fungal proteins in their respective primary amino acid sequences. The highly homologous protein sequences between the human host and the microbes suggest a reasonable probability that the autoantibodies in autoimmune diseases are derived from the host immunity against the microbes present in the human body, commensal or pathogenic, bacterial or fungal in origins. It is now known that human host is colonized by trillions of commensal microbes and these commensal microbes are intimate components of human development after birth (3-5). Exposure to environmental microbes including bacteria and fungi help develop normal immunity to prevent pathogenic infections, regulate human metabolic activity and various tissue functions (3-5).

It is noteworthy that no viral proteins are examined for autoantigens as the viral immunity is vastly different from the bacterial or fungal immunity. Anti-viral antibodies are protective against the subsequent viral infection, forming the basis of modern medical vaccination. However, exceptions are well-known that the viruses can evade the human immune system by hiding in the intracellular compartments and reactivated in response to the body stress conditions or immune compromised conditions. The examples of reactivation of Epstein-Bar virus (EBV) and herpes simplex virus in immune compromised patients are well-documented, although the mechanism of reactivation is yet to be proven. Specific anti-viral antibodies with cross reactivity to human tissues remain to be a possible mechanism for autoimmune diseases.

Production of antibody in vivo in response to a specific antigen is well studied, and a large scale production of antibody for pharmaceutical industry is performed routinely. However, the mechanism of the production of specific autoantibody in vivo, such as anti-CCP antibody (anti-CCP, cyclic citrullinated peptide, ACPA) in rheumatoid arthritis (RA) is still intriguing. Citrullination or deimination of protein is to convert the aiming acid arginine to amino acids citrulline. Citrulline is an unusual amino acid and it is not one of the standard 20 amino acids encoded by DNA in the genetic code. Citrullination is the result of post-translational modification by a group of enzymes arginine deiminases or peptidylarginine deiminase (PADs). Deimination of proteins will change the hydrophobicity since arginine is positively charged and citrulline is not, and the protein Citrullination leads to abnormal folding of proteins and its functions. Citrullination of proteins will also induce abnormal immune response by generating anti-citrullinated protein antibodies, leading to autoimmune disease such as RA and multiple sclerosis (MS). There are multiple PADs in human with enzymatic deiminase activities. Human PADs distribute in a variety of tissues in a tissue specific fashion, and these enzymes likely play important roles in cellular functions and signal transduction. There are also many bacterial deiminases that share significant homologies with human proteins. Bacterial deiminases, from either commensal or pathogenic, can potentially citrullinate human proteins, leading to abnormal citrullinated proteins to induce human immune response and autoimmune diseases. Alternatively, bacterial deiminases can directly induce human immune response upon entering the human body, and anti-deiminase antibody generated against the bacterial deiminases can cross-react to the human enzymes (deiminases), leading to autoantibodies and autoimmune diseases, although neither the reports of anti-deiminase autoantibodies in human diseases are documented nor any effort is made to discover anti-deiminase antibody in autoimmune diseases. However, in mouse model, a recent study demonstrated that the immunized mice with the murine or the human PAD2 or PAD4 enzyme can generate anti-citrullinated peptide anybody, and these antibodies against citrullinated peptides only occur with the enzyme (PAD) bound to the targets (14). It appears that the immunogenic epitope is the peptide-enzyme complex. Furthermore, the bacterial PAD from porphyromonase gingivalis (periodontitis) is important in periodontal disease and it is also important for RA (15). At this point, the data to support the presence of anti-bacterial PAD antibodies or anti- PAD autoantibodies in autoimmune diseases are lacking, although it is plausible with a new mechanism of disease process.

The present study of the autoantigens and the microbial proteins is a computational prediction and requires vigorous experimental validation. Specific epitopes eliciting antibody productions in vivo are usually small peptides, and BLASTP analysis could miss some small areas of amino acids that are antigenic. Antigenicity of any given protein is not random. Epitope mapping of human autoantigens may help answer these questions. Combination of direct comparison of the human autoantigens and the microbial proteins with specific antigenicity mapping can help anticipate the epitopes of any given proteins and facilitate the specific epitope mapping. These computational tools are widely available in public and make it possible to study the specific autoimmune diseases for better diagnostics and therapeutics. Many microbial proteins identified by the BLASTP with homology to the human autoantigens are hypothetical proteins from the known or unknown microbes, and significant effort is required to characterize these unknown microbes (bacteria or fungi) and the functions of their hypothetical proteins so that clinical significance of these microbial proteins can be determined. Some hypothetical proteins are derived from the well-known common microorganisms such as *E. coli*, *Enterobacters* or *Candida albicans*, and the functions of these microbes within the body and disease state is yet to be completely understood.

## Conclusion

More than two third of the autoantigens important for human autoimmune diseases are homologous to the microbial proteins, suggesting a majority of the autoantibody production are against the microbial protein origin. Further experimental validation is required for better understanding of the autoantibodies and the autoimmune diseases.

## References

1. Kumar V, Abbas, AK., Aster, JC. Robbins and Cotran Pathologic Basis of Disease - 9th edition. 9th Ed. ed.: Elsevier; 2015.

2. Kasper D, Fauci, AS., Hauser, SL., Longo, DL., Jameson, JL., Loscalzo, J. Harrison’s Principles of Internal Medicine. 19th Ed. ed.: McGraw Hill Education; 2015.

3. Gilbert JA, Blaser MJ, Caporaso JG, Jansson JK, Lynch SV, Knight R. Current understanding of the human microbiome. Nat Med. 2018 Apr 10;24(4):392–400.

4. Gilbert JA, Quinn RA, Debelius J, Xu ZZ, Morton J, Garg N, et al. Microbiome-wide association studies link dynamic microbial consortia to disease. Nature. 2016 Jul 7;535(7610):94–103.

5. Young VB. The role of the microbiome in human health and disease: an introduction for clinicians. Bmj. 2017 Mar 15;356:j831.

6. Kowarsky M, Camunas-Soler J, Kertesz M, De Vlaminck I, Koh W, Pan W, et al. Numerous uncharacterized and highly divergent microbes which colonize humans are revealed by circulating cell-free DNA. Proc Natl Acad Sci U S A. 2017 Sep 05;114(36):9623–8.

7. Zhang P, Minardi LM, Kuenstner JT, Zekan S, Kruzelock R. Extracellular Components in Culture Media of Mycobacterium Avium Subspecies and Staphylococci with Implications for Clinical Microbiology and Blood Culture. American Journal of Infectious Diseases and Microbiology. 2016;4(6):112–7.

8. Zhang P, Minardi, LM., Kuenstner, JT., Zekan, SM., Zhu, F., Hu, YL., Kruzelock, R. Cross–reactivity of antibodies against microbial proteins to human tissues as basis of Crohn’s disease and Sjogren’s syndrome. http://biorxivorg/content/early/2017/03/13/116574.2017.

9. Zhang P, Minardi LM, Kuenstner JT, Zekan SM, Kruzelock R. Anti-microbial Antibodies, Host Immunity, and Autoimmune Disease. Front Med (Lausanne). 2018;5:153.

10. Somayaji R, Priyantha MA, Rubin JE, Church D. Human infections due to Staphylococcus pseudintermedius, an emerging zoonosis of canine origin: report of 24 cases. Diagn Microbiol Infect Dis. 2016 Aug;85(4):471–6.

11. Kochibe N, Nicolotti RA, Davie JM, Kinsky SC. Stimulation and inhibition of anti-hapten responses in guinea pigs immunized with hybrid liposomes. Proc Natl Acad Sci U S A. 1975 Nov;72(11):4582–6.

12. Niegowska M, Paccagnini D, Burrai C, Palermo M, Sechi LA. Antibodies against Proinsulin and Homologous MAP Epitopes Are Detectable in Hashimoto’s Thyroiditis Sardinian Patients, an Additional Link of Association. PLoS One. 2015;10(7):e0133497.

13. Aschermann S, Lehmann CH, Mihai S, Schett G, Dudziak D, Nimmerjahn F. B cells are critical for autoimmune pathology in Scurfy mice. Proc Natl Acad Sci U S A. 2013 Nov 19;110(47):19042–7.

14. Arnoux F, Mariot C, Peen E, Lambert NC, Balandraud N, Roudier J, et al. Peptidyl arginine deiminase immunization induces anticitrullinated protein antibodies in mice with particular MHC types. Proc Natl Acad Sci U S A. 2017 Nov 21;114(47):E10169–E77.

15. Gabarrini G, de Smit M, Westra J, Brouwer E, Vissink A, Zhou K, et al. The peptidylarginine deiminase gene is a conserved feature of Porphyromonas gingivalis. Sci Rep. 2015 Sep 25;5:13936.

